# Dietary intervention in captive-bred hares fails to enrich gut microbiomes with wild-like functions

**DOI:** 10.1101/2024.12.03.626655

**Authors:** Ostaizka Aizpurua, Garazi Martin-Bideguren, Nanna Gaun, Antton Alberdi

## Abstract

Reintroducing captive-bred animals into the wild often faces limited success, with the underlying causes frequently unclear. One emerging hypothesis is that maladapted gut microbiota may play a significant role in these challenges. To investigate this possibility, we employed genome-resolved metagenomics to analyse the taxonomic and functional differences in the gut microbiota of wild and captive European hares (*Lepus europaeus*), as well as to assess the impact of a dietary switch to grass aimed at pre-adapting captive hares to wild conditions. Our analyses recovered 860 metagenome-assembled genomes, with 87% of them representing novel species. We found significant taxonomic and functional differences between the gut microbiota of wild and captive hares, notably the absence of Spirochaetota in captive animals and differences in amino acid and sugar degradation capacities. While the dietary switch to grass induced some minor changes in the gut microbiota, it did not result in a shift towards a more wild-like microbial community. The increased capacity for degrading amino acids and specific sugars observed in wild hares suggest that, instead of bulk grass, dietary interventions tailored to their specific dietary preferences might be necessary for pre-adapting hare gut microbiota to wild conditions.

**Importance:** This study sheds light on the critical role of gut microbiota in the success of reintroducing captive-bred animals into the wild. By comparing the gut microbiota of wild and captive European hares, we identified significant taxonomic and functional differences, including the absence of key microbial groups in captive hares. Dietary interventions, such as switching to grass, showed limited success in restoring a wild-like microbiota, highlighting the need for tailored approaches to mimic natural diets. With 87% of recovered microbial genomes representing novel species, this research also enriches our understanding of microbial diversity in wildlife. These findings emphasise that maladapted gut microbiota may hinder the survival and adaptation of reintroduced animals, suggesting that microbiome-targeted strategies could improve conservation efforts and the success of animal rewilding programs.

## Introduction

Amid the ongoing biodiversity crisis, captive breeding and release programs have become vital components of global conservation efforts (Gundu and Adia, 2014; Kleiman, 1989). These initiatives not only provide safe havens for threatened species but also create genetically diverse source populations for reintroduction into their natural habitats (Weeks et al., 2015). Despite big efforts to adjust captivity management practices to each species’ needs, the success rate of many reintroduction programmes remains limited, with the underlying causes often unclear (Jule et al., 2008; Seddon et al., 2007).

Although managers strive to replicate natural habitats as closely as possible, captive-bred animals experience lifestyles that differ significantly from those of their wild counterparts (Newberry, 1995; Young, 2013). When animals are released into the wild, they face drastic changes in diet, environmental exposure, foraging behaviour and social structures. To mitigate these transitions, many programs incorporate buffering strategies, such as exposure to a wild-like diet or confinement within natural habitats, which need to be tailored to the adaptability of each species (Laule, 2003; Shepherdson, 1994). Traditionally, the adaptive capacity of animals has been defined as a function of physiological and behavioural traits largely determined by the host genome (Frankham, 2008; Jamieson, 2011). Recent studies, however, have highlighted the critical role of gut microorganisms in many aspects of animal biology, including nutrition, immune function and behaviour (Lee and Hase, 2014; Park et al., 2018; Sherwin et al., 2019), suggesting that gut microbiota may significantly influence the adaptive capacity of animals reintroduced into the wild (Dallas and Warne, 2022; Diaz and Reese, 2021; West et al., 2019).

Research comparing the gut microbiota of captive animals with their wild counterparts has consistently revealed that captivity alters the gut microbiota (Allan et al., 2018; Bensch et al., 2023; Gibson et al., 2019; Guo et al., 2019; Koziol et al., 2022; Nakamura et al., 2011). However, the specific responses varied across species (Alberdi et al., 2021; McKenzie et al., 2017), indicating that a one-size-fits-all approach to managing animal health through microbiome modulation is unlikely to be effective. Although the exact mechanisms behind these differences are not fully understood, diet is considered a key factor. Due to management constraints, captive animals are often fed simplified dietary formulations, such as chow or pellets (Gibson et al., 2019; McKenzie et al., 2017; Mulder et al., 2016). Although these diets are nutritionally complete, they lack the structural complexity and indigestible components found in wild diets. These components are crucial in shaping the gut microbiota, particularly in hindgut fermenting herbivorous animals, as they are necessary for the production of essential fermentation metabolites like short-chain fatty acids, which have a direct impact on host health (Fang et al., 2020; Rechkemmer et al., 1988). As a result, maintaining, or gradually transitioning, captive animals to a more natural diet, has been proposed as a strategy to enhance their well-being and improve the success of reintroduction into the wild.

One of the reasons why the mechanisms behind the varying microbiota are poorly understood is that most prior studies have relied on 16S rRNA amplicon sequencing (Martínez-Mota et al., 2020; Padula et al., 2021; van Leeuwen et al., 2020), a method that, while useful, does not provide direct insights into the functional differences between microbial communities in captive versus wild animals. In contrast, genome-resolved metagenomics, which reconstructs bacterial genomes from metagenomic samples, enables the recovery of the functional capacities of each member of the gut microbiota (Scholz et al., 2012). This approach allows for a more detailed understanding of the metabolic contributions microorganisms can offer to their host, thus providing a much deeper resolution on the mechanistic processes behind microbial turnover between captive and wild populations.

In this study, we used the European hare (*Lepus europaeus* Pallas, 1778) as a study system to analyse the functional differences in gut microbiota between wild and captive animals using genome-resolved metagenomics. Additionally, we tested whether switching the diet of captive hares to grass could shift the functional capacities of their microbial communities to more closely resemble those of their wild counterparts. By doing so, we aim to better understand the role of gut microbiota in the success of reintroduction programs and enhance conservation efforts for the European hare.

## Results

### A genome catalogue of the European hare

We generated a total of 222.06 Gbp of sequencing data from 45 samples, with an average depth of 4.93 ± 1.08 Gbp per sample. The co-assembly and binning of the metagenomic data yielded 860 metagenome-assembled genomes (MAGs) (Figure 1A), with an average completeness of 80.99 ± 16.79% and contamination of 2.21 ± 2.32% (Figure S1). The MAG catalogue was dominated by Bacillota A (61%) genomes, followed by Bacteroidota (14.77%), with the rest of the phyla accounting for less than 10% of the genomes. Notably, 79 MAGs were not annotated at the genus level, and 749 MAGs (87.09%) lacked species-level annotations that differ depending on the phylum, from 33.85% (Bacteroidetes) to 100% (Actinomycetota, Patescibacteria, Desulfobacterota, Bacillota B, Campylobacterota) (Figure 1A). Overall, Bacillota A had the highest number of MAGs without species-level annotation, accounting for 516 MAGs (98.1%). MAGs exhibited a genome size range spanning 0.6 Mb (*Nanosyncoccus*, Patescibacteria) and 9.9 Mb (*Lawsonibacter*, Bacteroidota). The dereplicated MAG catalogue contained 1,362,707 redundant genes, out of which 1,045,555 (76.7%) received some annotation and 610,762 (44.8%) were assigned KEGG orthologs. MAGs exhibited a wide range of functional attributes, with Bacillota (A, B, C) showing the widest functional capacities (Figure 1B). MAGs belonging to the same phylum clustered together, suggesting that MAGs that are phylogenetically closer have higher similar functional attributes.

**Figure 1.**
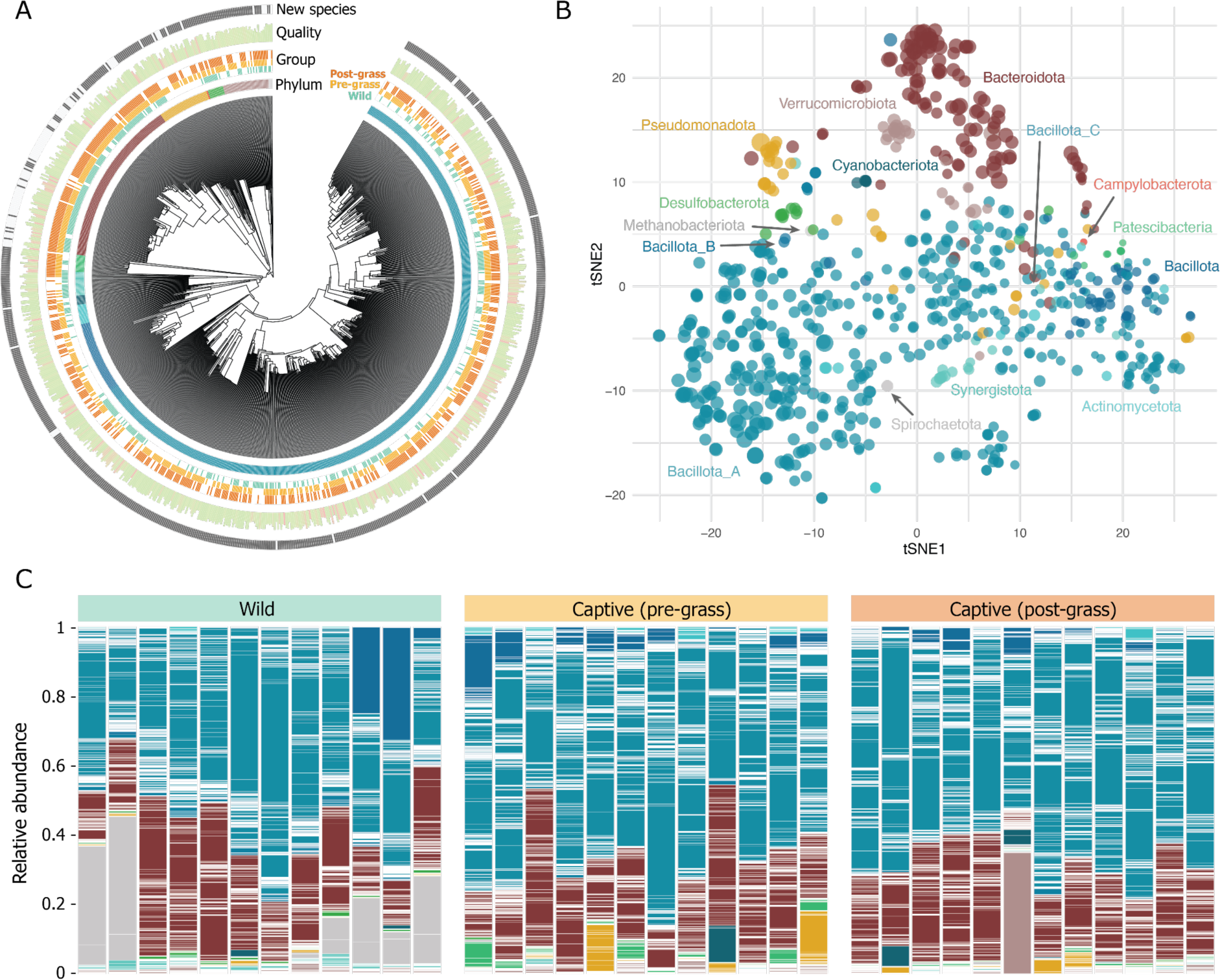
Overview of the metagenome-assembled genome (MAG) catalogue. **a)** Circular phylogenetic tree of the 860 MAGs reconstructed in this study. The first ring displays the phylum of the genome. The second ring indicates the quality of the genome, with the length of the bars showing completeness and colour indicating a contamination gradient from 0 (green) to the maximum value of 10 (red). The third ring indicates whether the genome is a new species (dark) or related (>95% average nucleotide identity) to a known species. **b)** tSNE ordination of the functional attributes of the genomes, coloured by phylum, with size indicating genome length. **c)** Relative abundances of the MAGs in samples collected in the wild and captive animals before and after grass feeding.

When the reads were mapped to the host genome, we observed that some wild samples contained a significantly higher fraction of host DNA, which caused an underrepresentation of microbial data. To address this, samples with a high host DNA fraction (n=9) were excluded from community analyses (Figure S2). This exclusion increased the average values and decreased the dispersion of domain-adjusted mapping rates (DAMR), from 86.5 ± 27.77% to 100%, thus increasing comparability across samples.

### Taxonomic and functional composition of wild and captive hares

The gut microbiota of European hares revealed the presence of 14 bacterial phyla, with Bacillota A (57.24 ± 12.62%) and Bacteroidota (25.79 ± 10.11%) being the most dominant across all samples (Figure 1C). These 14 bacterial phyla were consistently observed in both wild and captive-bred individuals, although significant differences in their relative abundances were noted (Table S1). While Bacillota A and Bacteroidota were the two dominant phyla in both groups, eight phyla were significantly different (Spirochaetota, Bacillota B, Synergistota, Bacillota C, Desulfobacterota, Verrucomicrobiota, Bacteroidota, Bacillota A, p<0.05). For example, wild hares exhibited a notable presence of Spirochaetota (11.73 ± 14.79%), which was nearly absent in captive individuals (0.00063 ± 0.0007%). Additionally, Archaea, specifically Methanobacteriota, were detected exclusively in captive-bred individuals.

Further analysis revealed six MAGs from Pseudomonadota (present in 55.56 ± 16.39% of the samples), one MAG from Bacillota A (50%), and one Archaea MAG from Methanobacteriota (66.67%) to be unique to the captive-bred hares. One MAG was found to be present only in wild individuals (Pseudomonadota), but it only appeared in two individuals. Differential abundance analysis identified 342 MAGs representing 116 genera that were significantly different between wild and captive-bred hares. Taxonomic diversity analyses revealed significant differences in both alpha and beta diversities between wild and captive-bred hares. Specifically, alpha diversity analyses indicated statistically significant differences in richness and neutral diversity (Figure 2A), though phylogenetic diversity did not show significant variation. All metrics were statistically significant in beta diversity, with neutral diversity (R^2^=0.254, p=0.001, Figure 2B) and phylogenetic diversity (R^2^=0.308, p=0.001) showing clear distinctions between wild and captive-bred hares.

**Figure 2.**
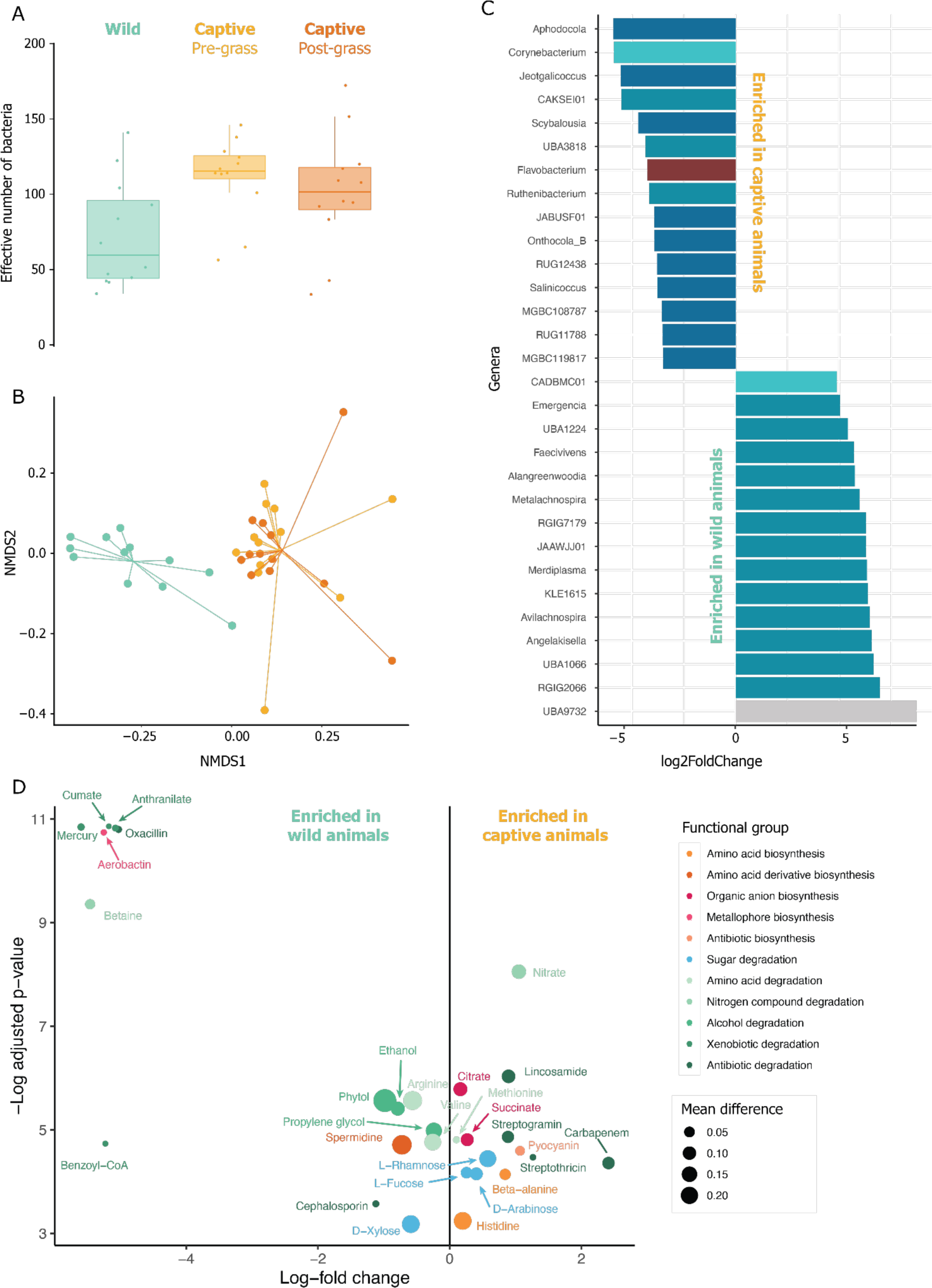
Diversity, taxonomic and functional microbiome differences between wild and captive hares. **a)** Alpha diversity differences between the three analysed groups in terms of Hill numbers of q=1 (exponential of Shannon index). **b)** Beta diversity differences between the three analysed groups in terms of Hill numbers of q=1. **c)** Differential abundance of MAGs between wild- and captive-born animals, in which MAGs with significant log-fold differences between sample types are coloured according to their phyla. **d)** Differential abundance of functions between wild- and captive-bred animals.

In terms of gut microbiota functional capacities, functional alpha diversity was found to be statistically significant (t = 4.3493, df = 21.934, p-value = 0.00026), whereas functional beta diversity was not (R^2^=0.188, p=0.086). The Metabolic Capacity Indices values did not differ significantly between the two groups (0.249±0.0152 in captive-bred vs. 0.254±0.0206 in wild individuals; t = −0.65356, df = 20.27, p-value = 0.521). However, 14 functions were enriched in captive individuals, while 15 were enriched in wild individuals (Figure 2C). In captive hares pathways for four antibiotics (Streptothricin, Carbapenem, Lincosamide and Streptogramin), an amino acid (Methionine), a nitrogen compound (Nitrate), and two sugar (D-Arabinose, L-Rhamnose and L-Fucose) degradations were enriched, as well as the biosynthesis of two organic anions biosynthesis (Citrate, Succinate) and two amino acids biosynthesis (Beta-alanine and Histidine), and an antibiotic pyocyanin. On the contrary, wild hares possessed gut bacteria with a higher capacity for degrading amino acids (Valine, Arginine) and synthesising amino acids derivative spermidine compared to their captive-bred counterparts. Pathways for the degradation of four xenobiotics (Benzoyl-Coa, Mercury, Cumate, Anthranilate), three alcohols (Propylene glycol, Phytol, Ethanol), two antibiotics (Cephalosporin, Oxacillin), nitrogen compounds (Betaine), D-xylose sugar as well as metallophore aerobactin biosynthesis were also enriched in wild hares.

### Effect of dietary switch to grass in captive hares’ microbiota

To determine if the introduction of grass into the diet could pre-adapt the gut microbiota of European hares and enhance their adaptation capacity, we first assessed whether this dietary change produced any alterations in microbial composition and function in captive-bred animals. Our analysis revealed no significant differences in alpha (Figure S3) and beta (Figure S4) diversity in the individuals before and after grass was included in the diet (p<0.05). However, differential abundance analysis identified 11 Metagenome-Assembled Genomes (MAGs) that differed between the groups (Figure S5). Of these, eight MAGs were enriched in hares on a diet without grass, while three MAGs increased in abundance following the inclusion of grass in the diet. Specifically, the introduction of grass resulted in the increased presence of two MAGs from the Lachnospiraceae and Oscillospiraceae families within the Bacillota A phylum, and one MAG from the Xanthomonadaceae family within the Pseudomonadota phylum. In terms of functional differences, only two traits showed significant variation between the groups. The abundance of bacteria with the capacity to biosynthesize the metallophore staphylopine and degrade the antibiotic chloramphenicol decreased when grass was added to the diet.

## Discussion

Our metagenomic analyses of wild and captive European hares (*Lepus europaeus*) represent the first effort to reconstruct the species’ gut microbiota using genome-resolved metagenomics. We successfully recovered 860 genomes, 87% of which were previously unknown, which is in line with other wildlife microbiome analyses that also reported novel species rates over 75% (Leonard et al., 2024; Levin et al., 2021). This biodiscovery rate reflects the vast unexplored diversity within microbial communities associated with vertebrates, underscoring the importance of using shotgun metagenomics not only for detecting new taxa but also to uncover the distribution of microbial functions. In line with previous studies (Padula et al., 2021), the gut microbiota of hares was highly dominated by bacteria from Bacillota A and Bacteroidota, suggesting a role in breaking down complex polysaccharides from plant fibres and producing short-chain fatty acids as a result of their fermentation.

### Differences between captive and wild hares

Our analyses unveiled significant differences in the taxonomic composition and functional capabilities of the gut microbiota between captive and wild European hares, a trend that has been observed across various vertebrate species (Alberdi et al., 2021; Bensch et al., 2023; Feng et al., 2024; Liu et al., 2021, 2024; McKenzie et al., 2017). The transition to or breeding within a captive environment usually leads to shifts in microbial communities, which appear to be largely influenced by species-specific and environmental conditions within captivity (Alberdi et al., 2021). However, it remains uncertain whether these changes result from dysbiosis, represent adaptations to the captive setting or are neutral towards the host. Captive and wild hares in this study shared the same 14 bacterial phyla, with Bacillota and Bacteroidota being dominant in both groups, consistent with prior findings in lagomorphs (Stalder et al., 2019; Velasco-Galilea et al., 2018; Zhao et al., 2024). However, captive hares had higher neutral alpha diversity than their wild counterparts, indicating that captivity introduced a variety of additional microbial taxa. Among these, seven bacterial species were unique to captive hares, comprising six Pseudomonadota (five MAGs from Gammaproteobacteria and one from Alphaproteobacteria) and one Bacillota A (Clostridia). Additionally, the only archaeal genome reconstructed, *Methanosphaera cuniculi*, was also exclusive to captive hares. *Methanosphaera* has been identified in other animals such as squirrels (Carey et al., 2013), rabbits (Kušar and Avguštin, 2010), pigs (Luo et al., 2012; Peng et al., 2022), humans (Mohammadzadeh et al., 2022), and ruminants (Smith et al., 2022), highlighting its adaptability to different host gut environments. Notably, *Methanosphaera* exhibits host-specificity, with reduced genome sizes in monogastric hosts (∼1.7 Mbp) compared to larger genomes found in ruminants (∼2.9 Mbp) (Borrel et al., 2020). The genome size of the reconstructed *Methanosphaera* in captive hares (1.7084 Mbp) closely resembled those found in monogastric animals (Borrel et al., 2020). Although these archaea are well-established contributors to fibre fermentation in ruminants and other monogastric herbivores, their role in hares remains less defined. These archaea can contribute to hydrogen turnover in the gut, thereby potentially aiding fermentation efficiency. Yet, in our study, *Methanosphaera* was not present in wild hares, and its absence did not appear to affect polysaccharide degradation capacities, suggesting that it may not be essential for fibre fermentation in hares. This finding contrasts with studies in ruminants, where methanogens play a more central role in fermentation converting hydrogen into methane to maintain balance in the gut environment (Hook et al., 2010). The lack of methanogens in wild hares, alongside comparable microbial degradation capacities between wild and captive animals, supports the view that archaea may not be necessary for optimal fermentation in monogastric herbivores like hares in natural settings. However, their potential significance in captivity remains an open question for further investigation.

The relative abundances of several bacteria differed significantly between captive and wild hares, with the most notable difference observed in the phylum Spirochaetota. This phylum has previously been reported as a prominent component of hare microbiota, with a larger representation compared to rabbits (Stalder et al., 2019). The two MAGs identified in this study lacked species-level annotation but were classified within the genus *UBA9732* (Sphaerochaetaceae). Despite being the third most abundant phylum in wild hares, Spirochaetota were nearly absent in captive-bred individuals. This absence could be attributed to several factors, including the unavailability of these bacteria in captivity or the younger age of the captive hares, as Spirochaetota may colonise hare guts later in life. However, a prior study also reported no Spirochaetota among the top genera in captive hares (Padula et al., 2021), reinforcing the idea that limited exposure to these bacteria in captivity may be a contributing factor.

Interestingly, *UBA9732* has also been found in the rumen of herbivores (Hernández et al., 2022) and in the gut microbiomes of populations with high fibre intake, such as South African (Tamburini et al., 2022) and Baka forager-horticulturalists from Cameroon (Rampelli et al., 2024). The enrichment of Spirochaetota has been observed in rural populations (Angelakis et al., 2019; De Filippo et al., 2010) while a significant decline in these bacteria has been noted in westernised humans and platyrrhines, likely due to lifestyle changes (Thingholm et al., 2021). Our findings suggest that the limited presence of these anaerobic, non-spore-forming bacteria in captive hares is likely a result of altered diets and reduced social interaction in captivity (Thingholm et al., 2021). The two Spirochaetota genomes reconstructed in this study exhibited strong capacities for degrading alpha- and beta-galactans, as well as arabinan, which could enhance the digestion of complex plant fibres in the lower intestines of hares. This ability may be particularly advantageous for wild hares consuming fibre-rich diets, but less relevant for captive hares on controlled, processed diets.

Overall, the gut microbial communities of wild and captive hares displayed notable differences in their functional capacities. Wild hares exhibited enhanced microbial abilities to degrade D-xylose, while captive-bred hares showed enriched capacities for degrading L-rhamnose and L-fucose. D-xylose is abundant in hemicellulose, a major component of plant cell walls, particularly in hardwoods and grasses. In contrast, L-rhamnose and L-fucose are abundant in softer plants. The distinct sugar degradation profiles observed between wild and captive hares probably reflect their adaptation to diets with distinct sources of carbohydrates.

Similarly, wild hares were found to host gut microbiota with greater capacities for amino acid degradation compared to their captive counterparts. This enhanced microbial function may also be linked to the dietary preferences of hares, particularly during autumn and winter, when they tend to favour plants that are low in fibre but rich in crude fat and protein (Schai-Braun et al., 2015). These seasonal dietary shifts could select gut microorganisms with an increased ability to break down amino acids, providing hares with essential nutrients during periods when high-protein foods are more abundant. This functional adaptation in their microbiota likely helps wild hares maximise nutrient extraction from their protein-rich diet, supporting their survival in changing environmental conditions. In contrast, amino acid biosynthesis was found to be higher in captive hares, suggesting that their gut microbiota might compensate for a diet potentially lower in protein by producing essential amino acids. This increased biosynthetic capacity may reflect a more controlled and less varied diet in captivity, where external protein sources are limited. As a result, captive hares presumably have to rely more on their gut microbes to source amino acids, ensuring they meet their nutritional requirements despite a less protein-rich diet. These differences in amino acid metabolism highlight the influence of diet on microbial functions and the adaptive responses of gut microbiota in different environments.

Furthermore, the gut microbial community of wild hares exhibited a higher capacity to degrade xenobiotics, although the mean difference between wild and captive hares was relatively small. The presence of bacteria with the ability to break down xenobiotics might be because of the exposure of wild hares to these environmental contaminants (De Filippis et al., 2024) or indirectly, it turns out that the necessary bacteria found in wild hares have these capabilities as well. In natural environments, animals encounter various xenobiotics, including pollutants and plant- or industry-derived chemicals. Future studies could explore whether this microbial capacity for metabolising xenobiotics functions as a protective adaptation (Claus et al., 2016), helping wild hares detoxify and process harmful substances like mercury, which pose serious risks to animal health (Gworek et al., 2020).

### Effect of dietary switch to grass in captive hares

Gut microbiota has been proposed as a tool to pre-adapt captive-bred animals to wild conditions, potentially improving the success of reintroduction programs (Mulder et al., 2016; van Leeuwen et al., 2020). In this study, we examined whether switching captive-bred hares from a pellet-based diet to grass could modify their gut microbial community, aligning it more closely with that of wild hares. Contrary to a previous study (Padula et al., 2021), we observed that this dietary change led to only minimal alterations in the gut microbiota. For instance, we notice a decrease in *Eggerthellaceae* after the introduction of grass. This bacterial family is associated with a chow diet in mice, likely due to its high content of cereal-derived fibres (Rodríguez-Daza et al., 2020). We also observed an increase in the relative abundance of *Luteimonas* following the dietary switch. This genus has been detected in the phyllosphere of various plants (Wemheuer et al., 2017), suggesting that hares may have been exposed to this bacterium through their new food source. Similarly, the microbial community showed few functional changes. The decrease in bacteria that produce staphylopine after grass introduction could potentially decrease the infection risk of the animals, improving animals’ health in captivity. Additionally, the observed loss of the capacity to degrade chloramphenicol may be linked to the decrease in *Eggerthellaceae*, a group known for its high capacity to break it down. While relative abundances of certain bacteria differed, the inclusion of grass in captivity did not include new bacteria species nor significantly enhance the functional capacities of hares in a way that would pre-adapt them to the wild environment.

More importantly, the dietary switch to grass did not make the gut microbiota of captive hares more similar to that of wild hares. The differences in fibre and amino acid degradation capacities between captive and wild hares suggest that using specific plant sources, rather than generic grass, might be crucial in aligning captive microbiomes with those of wild hares. A detailed analysis of wild hares’ dietary habits could help identify protein-rich plants that contribute to the functional differences between wild and captive animals. Incorporating these plants into the diet of captive hares could serve as a more effective pre-adaptation strategy for reintroduction, providing a better alternative to grass.

## Conclusion

This study demonstrated the value of combining genome-resolved metagenomics with functional microbiome analyses to uncover key functional differences in the gut microbial communities of wild and captive European hares. The finding that simply introducing grass into the diet does not sufficiently align the microbiomes of captive hares with those of wild individuals, alongside the functional discrepancies observed, suggest that successful animal pre-adaptation strategies require a more nuanced understanding of species-specific dietary habits, beyond superficial dietary changes. To better prepare captive animals for reintroduction into the wild, it will be essential to develop diets that not only mimic natural plant compositions but also support the microbial functions that are critical to overcoming adaptation challenges they experience, such as diarrhoea, in natural environments. By tailoring reintroduction programs to optimise gut health and microbiome functionality, we may improve the resilience of captive-bred animals and enhance their chances of survival in the wild. As habitats continue to change and human impacts on ecosystems intensify, understanding how gut microbiota influences animal adaptation and health will be increasingly critical for the success of species conservation efforts.

## Material and Methods

### Captive-breeding programme

This research was conducted within the framework of the captive-breeding program of the Regional Government of Gipuzkoa (Basque Country), located at the southern edge of the European hare’s geographical distribution. This action aims to restock hare populations in the region, which have undergone a significant decline over the past four decades. The program involves a carefully managed reintroduction process, where captive-bred hares are gradually acclimatised to the wild environment. A key aspect of this process is the introduction of grass into their pellet-based diet (C1, Unamuno, Altsasua) before release into the natural habitat. Despite these efforts, some animals experience adaptation challenges, such as diarrhoea, after being released.

### Sampling

A total of 24 faecal samples were obtained from animals bred at the hare breeding facility in Altsasu, The Basque Country (42.88°, −2.19°). Faecal samples were collected twice from each individual: once before (n=12) and 10 days after grass (n=12) was introduced into their diet. In addition, samples from 21 wild hares were also collected leveraging animals captured by hunters in the region. Immediately after the animals were hunted, forest rangers from the Regional Government of Gipuzkoa collected faecal samples directly from the colon following an EHI standardised protocol (https://www.earthhologenome.org/sampling.html). All faecal samples were placed in tubes containing 1 ml of DNA/RNA Shield (Zymo, USA) and stored in a refrigerator before being shipped to the lab, where they were kept at −20°C until DNA extraction.

### Laboratory work

Sample processing was conducted following the EHI data generation procedures (www.earthhologenome.org/laboratory) (Leonard et al., 2024; Pietroni et al., 2024). In short, chemically digested samples were subject to mechanical cell lysis (bead-beating) using Tissuelyzer (Qiagen, USA) to release as much DNA as possible from different kinds of prokaryotic cells. Subsequently, DNA was isolated and purified using silica magnetic beads combined with solid-phase reversible immobilisation to remove as many inhibitors as possible. Extracted DNA was quantified using a Qubit 3 fluorometer (Thermo Fisher Scientific), and DNA quantities levelled to 200 ng of DNA in 24 μl of water before shearing the DNA using a Covaris LE220 ultrasonication device to adjust DNA fragment sizes to efficient short-read sequencing. Adapter ligation-based sequencing library preparation was then conducted using the in-house developed BEST protocol (Carøe et al., 2018). Library quality was screened through qPCR and gel electrophoresis before conducting library amplification with dual index identifiers with an adjusted number of PCR cycles for each library to minimise the number of artefacts and loss of library complexity derived from sample overamplification. Finally, uniquely indexed libraries were merged in multiple library pools at exact molarities to produce around 8GBs (gigabases) of 150 bp paired-end sequencing data per sample. Sequencing was carried out in multiple sequencing runs at an Illumina NovaSeq X platform by the sequencing provider Novogene (UK).

### Bioinformatics

Data processing was carried out using the standard EHI bioinformatics pipeline (www.earthhologenome.org/bioinformatics). The pipeline is based on snakemake (Köster and Rahmann, 2012) for workflow management, conda for environment management and slurm for computational job management, and it is directly managed from the EHI database. Data were first quality-filtered using fastp (Chen et al., 2018) before splitting the metagenomic from the host genomic fraction through mapping reads against the reference *Lepus europaeus* genome (GCF_033115175.1) using bowtie2 (Langmead and Salzberg, 2012). Subsequently, the metagenomic fraction underwent the genome-resolved metagenomic pipeline. Samples were both assembled individually and co-assembled at the origin level using Megahit (Li et al., 2015). Assembly contigs were clustered into bins using Maxbin2(Wu et al., 2016), Metabat2 (Kang et al., 2019) and CONCOCT (Alneberg et al., 2014), followed by bin refinement using Metawrap (Uritskiy et al., 2018), and bin quality assessment using CheckM2 (Chklovski et al., 2023). Only bins with completeness values over 50% and contamination values under 10% were considered for downstream analyses (Bowers et al., 2017). All resulting metagenome-assembled genomes (MAGs) were dereplicated using dRep (Olm et al., 2017), before annotating them taxonomically against the GTDB database using GTDB-tk (Chaumeil et al., 2022) and functionally against the Pfam, KEGG, UniProt, CAZY, and MEROPS databases using DRAM (Shaffer et al., 2020). Finally, quality-filtered reads from each sample were mapped against the annotated MAG catalogue using bowtie2 (Langmead and Salzberg, 2012) to quantify the representation of each MAG in each sample using CoverM. We used distillR (https://github.com/anttonalberdi/distillR) to convert functional annotations into Genome-Inferred Functional Traits (GIFTs). This package provides quantitative measures of each MAG’s ability to degrade or produce important biomolecules, using a reference database containing 328 metabolic pathways and modules sourced from the KEGG (Kanehisa et al., 2016) and MetaCyc (Caspi et al., 2018) databases, which enables the transformation of raw annotations into 190 distinct GIFTs.

### Data analysis

All statistical analyses were conducted using R software v.4.3.2 (R Core Team, 2023). We calculated the alpha diversities of microbial communities using Hill numbers (Hill, 1973). To capture the effects of different diversity components (neutral, phylogenetic, and functional) and diversity orders (q = 0 considers only presence/absence, while q = 1 gives weight to MAGs based on their relative abundances), we calculated species richness at q = 0, neutral diversity at q = 1, phylogenetic diversity at q = 1, and functional diversity at q = 1 using the *Hilldiv2* package v.2.0.2 (Alberdi and Gilbert, 2019). Alpha diversity differences were determined by parametric t-test when the data was normally distributed and when the variances of the two groups were equal. When this assumption was not held, a non-parametric Wilcoxon test was performed. Linear mixed models, as implemented in the *lme4:lmer* function (Bates et al., 2015), were employed to evaluate differences in bacterial alpha diversity between captive-bred individuals before and after the grass was included in the diet, with hare ID included as a random effect.

We calculated the MAG composition dissimilarities between different samples using Hill numbers by computing the Jaccard-type turnover for neutral, phylogenetic, and functional beta diversities at order q = 1 using *hilldiv::hillpair*. To visualise the variation in microbial composition, we performed nonmetric multidimensional scaling (NMDS) ordination plots based on the derived distance matrices. Differences in dispersion within sampling methods were assessed using the *betadisper* function in the *vegan* package (Oksanen et al., 2013). To test for differences in microbial composition between samples, we conducted a PERMANOVA using the *adonis2* function in vegan. When comparing captive-bred individual differences before and after the change in diet, hare ID was included as a blocking factor to control for repeated sampling using the *strata* function. Additionally, to visualise bacteria according to their functional traits, MAGs were ordinated based on their GIFTs (Genomic Information Functional Traits) through a t-SNE analysis using the *Rtsne* package v.0.17 (Krijthe et al., 2017). Metabolic Capacity Index (MCI) was calculated as a quantitative metric of the metabolic contributions gut microbiota can confer to their hosts using distillR.

We performed differential abundance analyses to identify microbial taxa that significantly differ between samples using *ANCOM-BC2* (Lin and Peddada, 2024). These analyses accounted for the random effect of hare ID when analysing repeated individuals before and after the dietary change. Differential abundance analyses were conducted at the MAG and phylum levels. Additionally, we calculated community-weighted values of GIFTs before comparing values between groups using the Wilcoxon test. The p-values were adjusted to account for multiple tests using the Bonferroni method.

## Acknowledgements

We would like to thank all the hunters, forest rangers from the Regional Government of Gipuzkoa and Leocadio Galán who participated in the fieldwork and provided the samples, as well as the support people who facilitated the administrative paperwork, with a special mention to Iñaki Olano, who coordinated the sample collection. Finally, we thank Prof. Tom Gilbert for supporting the study and for his valuable feedback on the manuscript. This work was supported by the Carlsberg Foundation through the grant CF20-0460 and the Danish National Research Foundation under the grant DNRF143 “A Center for Evolutionary Hologenomics”. The funders had no role in study design, data collection and interpretation, or the decision to submit the work for publication.

## Data accessibility

Raw sequencing data and microbial genome sequences have been or will be published as part of upcoming EHI Data Releases, under the umbrella Bioproject PRJEB51837. The data tables containing the quantitative information analysed and the R scripts used for statistical analyses are available in a dedicated GitHub repository (https://alberdilab.github.io/lepus_metagenomics).

